# *miRCurator*: A rapid curation and analysis tool for plant microRNA studies

**DOI:** 10.1101/2023.01.10.523394

**Authors:** Bihter Avşar, Danial Esmaeili Aliabadi

## Abstract

Plant microRNAs (miRNAs) are tiny, non-coding RNAs that regulate the biological pathways either by inducing translational repression or messenger RNA decay. The advent of sequencing technologies increases the number of biological tools and computational methods to identify miRNAs with their targets inside the genome and transcriptome. In this study, we present a rapid curation software tool, the so-called *miRCurator*, which filters out unwanted microRNA candidates based on three significant criteria: (1) the pre-miRNA sequence should be deprived of multi-branched loops; (2) less than four mismatches on between mature miRNA and miRNA* strand are allowed; (3) no mismatches at DICER cut points are permitted on a mature miRNA strand. *miRCurator* is a stand-alone user-friendly package that does not rely on cutting-edge computing platforms. We have examined our tool on different organisms, which are shown as case studies, and it was able to successfully identify all the unwanted miRNA candidates. The developed tool helps to identify genuine miRNAs and provides reliable outcomes in an automated fashion by eliminating false-positive predictions.

## 1. Introduction

RNA silencing is an evolutionarily conserved mechanism in eukaryotes, which assists plants to carry out different functional processes in the organism. This mechanism is induced by double-stranded RNA (dsRNA) or hairpin structured RNA (hpRNA), consisting of common elements including DICER or DICER-like (DCL), and Argonaute (AGO) family proteins. RNA silencing in plants can be classified into four distinct pathways: microRNA (miRNA) pathway, trans-acting small interfering RNA (tasiRNA) pathway, RNA-directed DNA methylation pathway, and exogenic RNA silencing pathway. Plants have evolved multiple RNA silencing factors for diverse RNA silencing pathways. For instance, the model plant Arabidopsis encodes four DCLs, six RNA-dependent RNA polymerases (RDRs), and ten AGOs, plus several other factors [1]. RNA silencing mechanism of the plants is also used in biotechnology to increase disease resistance, improve the commercial traits of the plants, change their flowering time, and develop high-value industrial products [1].

In this study, we mainly focus on the microRNA elements and their identification methodologies in the genome and transcriptome. Plant microRNAs are tiny (i.e., ~ 21 nucleotide) endogenous and non-coding RNAs, which regulate the gene expression either by inducing translational repression or messenger RNA decay. Plant miRNAs are generated from specific stem regions of single-stranded hairpin precursors, which is one of the distinct features from other small RNAs. The abundance of intracellular miRNA is regulated under multiple levels of control mechanism including transcription, processing, RNA modification, RNA-induced silencing complex (RISC) assembly, miRNA-target interaction, and turnover [2]. With the emergence of next-generation sequencing technologies, new computational methods and approaches are being developed to analyze the big data resulted from sequence information. These new methodologies provide better economy and more information accuracy to the users in a shorter computation time. Similar computational tools are also produced for plant miRNA identification from genomic and/or transcriptomic data, such as miRkwood [3], miRPlant [4], C-mii [5]. Some of these tools are available on the web to offer remote access to a broader range of users.

Other computational approaches benefit from the homology-based prediction for miRNA identification. The homology-based conserved approach is implemented in SUmiRFind and SUmiRFold Perl scripts [6]. For the homology-conservation method, the reference miRNA sequences should be selected from the current release of miRBase [7], which is the most detailed and updated database. SUmiRFind exploits BLASTN algorithm [8] with parameters, and records the important hits in a file. SUmiRFold script creates the secondary structure of these hit sequences by folding them using the UNAFold, an implementation of Zuker algorithm [9]. The output file from the SUmiRFold script is in a tabular format, which includes the following information for each putative miRNA: The new miRNA sequence, conserved miRNA sequence, pre-miRNA sequence, sequence ID, mature miRNA length, pre-miRNA length, number of mismatches to the query, pre-miRNA stem-loop start and end sites, hairpin location, MFE (Δ*Gkcal/mol*), %GC content and MFEI.

This output file should be re-checked and some of the candidates may be removed by the users manually since there might be some miRNA candidates in the resulted file that do not satisfy these criteria: (1) the pre-miRNA sequence should not consist of multibranched loops; (2) less than four mismatches on between mature miRNA and miRNA* strand are allowed [10]; (3) no mismatches at DICER cut points are permitted on the mature miRNA strand. Nonetheless, the output file may have thousands of miRNA candidates based on the size of the source file; therefore, a manual quality control procedure is time consuming for the users. To ease the process, we developed a script, the so-called *miRCurator*, that eliminates these improper miRNAs from the SUmiRFold output file. Our script is a stand-alone user-friendly package that rapidly omits false-positive miRNA candidates, and generates report in both tabular and visual formats. Our previous studies have been subjected to *miRCurator* as case studies for the curation of miRNAs from the kiwifruit genome [11], hazelnut genome [12,13], spinach transcriptome [14], fabaceae family members [15], and medicinal plant transcriptome [16,17].

## 2. Materials and Methods

In this section, we provide the detail of our script in two subsections: In Section 2.1, the requested data structure is explained, by which the information is sent to be processed. In Section 2.2, we explain the developed tool in detail.

### 2.1. Preparation

Before all the steps, SUmiRFind, SUmiRFold and *miRCurator*, contigs (assembled raw reads) should be prepared as inputs for the miRNA identification in trancriptomic data, whereas, in genomic data, raw reads may be used if they are longer than 200 nt. The methodology for miRNA identification using homology conservation approach is summarized in Figure 1.

**Figure 1.**
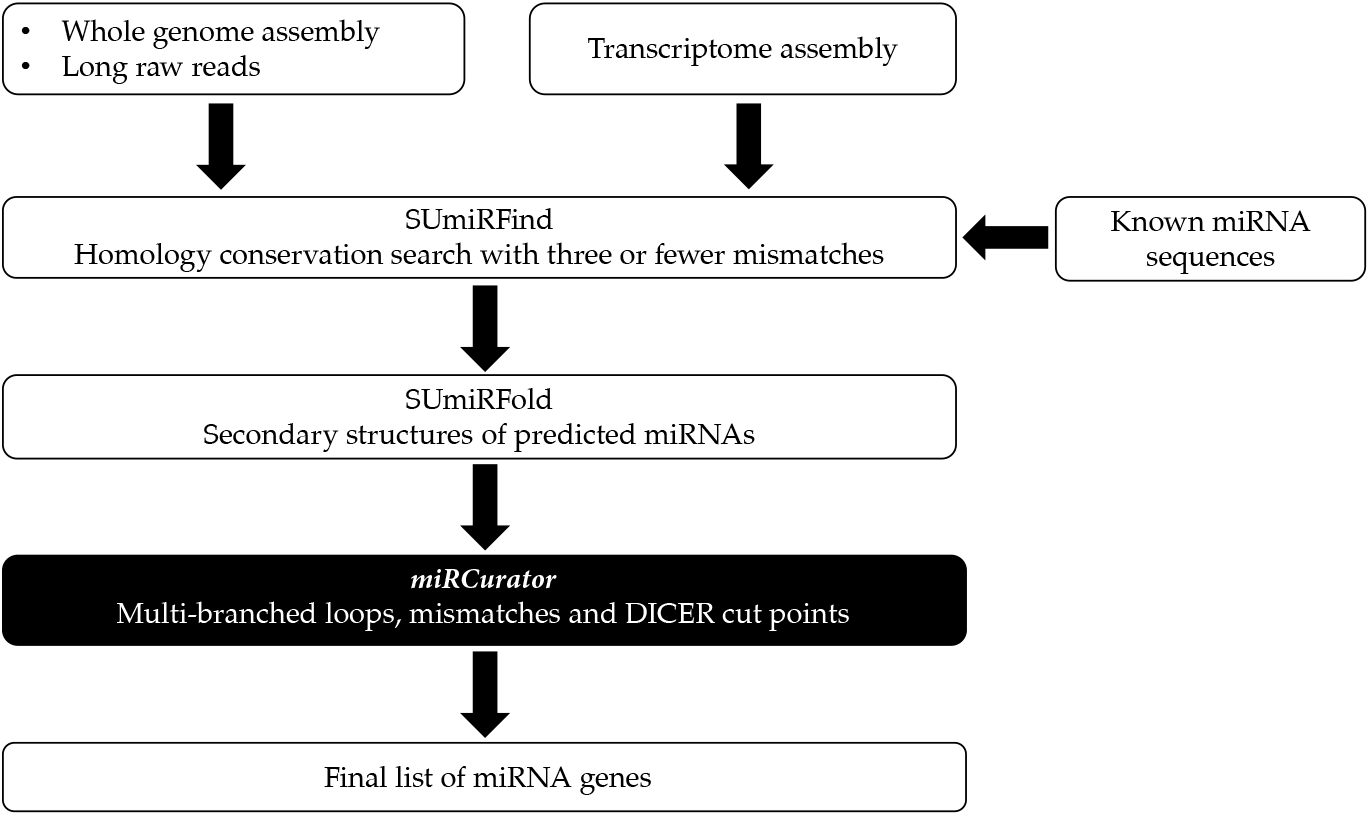
An overview of miRNA identification method.

As shown in Figure 1, the input data (e.g., whole genome assembly, long raw genomic reads, and transcriptome assembly) are used for SUmiRFind script in FASTA file format. In this step, the inputs align to known miRNA sequences with four or fewer mismatches. This alignment file is used for SUmiRFold part to search for the secondary structures. The SUmiRFold output files are provided in tabular format and they can be transformed to Microsoft Excel.

In order to prepare the inputs for *miRCurator*, users need to arrange the SUmiRFold output file on MS Excel with the following structure:

- The first column includes links to postscript files (*file_name.fsa_1.ps*). These addresses can also be used as a hyperlink to double-check results,
- the second and third columns are *Mature Start* and *Mature End*, respectively, and
- the fourth column is the *Unique Hit ID*.

### 2.2. miRCurator *Script*

After acquiring the data in MS Excel, the developed code (*miRCurator)* can be added as a Visual Basic for Applications (VBA) script. Since the VBA is embedded in Microsoft Excel, researchers can utilize the tool directly without further complication. The code can be divided into two main algorithms: (1) Multi-branched loop detector, and (2) Gene structure analyzer.

In order to conduct analysis on each gene, *miRCurator* reads each row of the Excel file and loads the corresponding *ct* file, which is created by the SUmirFold Perl script [6]. The associated gene in the Excel file is painted in yellow when the script cannot locate the *ct* file. In the generated *ct* file, the sequence of bases and their connections are mentioned. *miRCurator* reads the entire *ct* file and generates an *n_base_* × 8 matrix (**C**_*n*_*base*_×8_) where *n_base_* is the number of bases in the corresponding miRNA sequence. The columns in **C** matrix represents: (1) the index of current base in the miRNA sequence; (2) the corresponding base (i.e., A, C, G, and U); (3) the index of the previous base in the sequence; (4) the index of the next base in the sequence; (5) the index of the paired base; (6) the index of current base (repeated); (7 and 8) these columns specify whether the current base is paired (otherwise, zero values are associated to both for that row).

Algorithm 1 in *miRCurator* checks the sequence and counts the number of loops by looking at the 5th column of the **C** matrix. The time complexity of the Algorithm 1 is polynomial, in order of 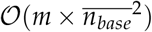 where *m* is the number of genes in the Excel file and 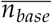 is the maximum sequence length in each gene. As displayed in Figure 2, the tip of each branch can be detected using the suggested algorithm. An advantage of this method is faster scan speed, which is due to ignoring long stems in the first loop (i.e., the condition over **C**_i,5_ in the 5th line).

#### Algorithm 1: Multi-branched loop detector

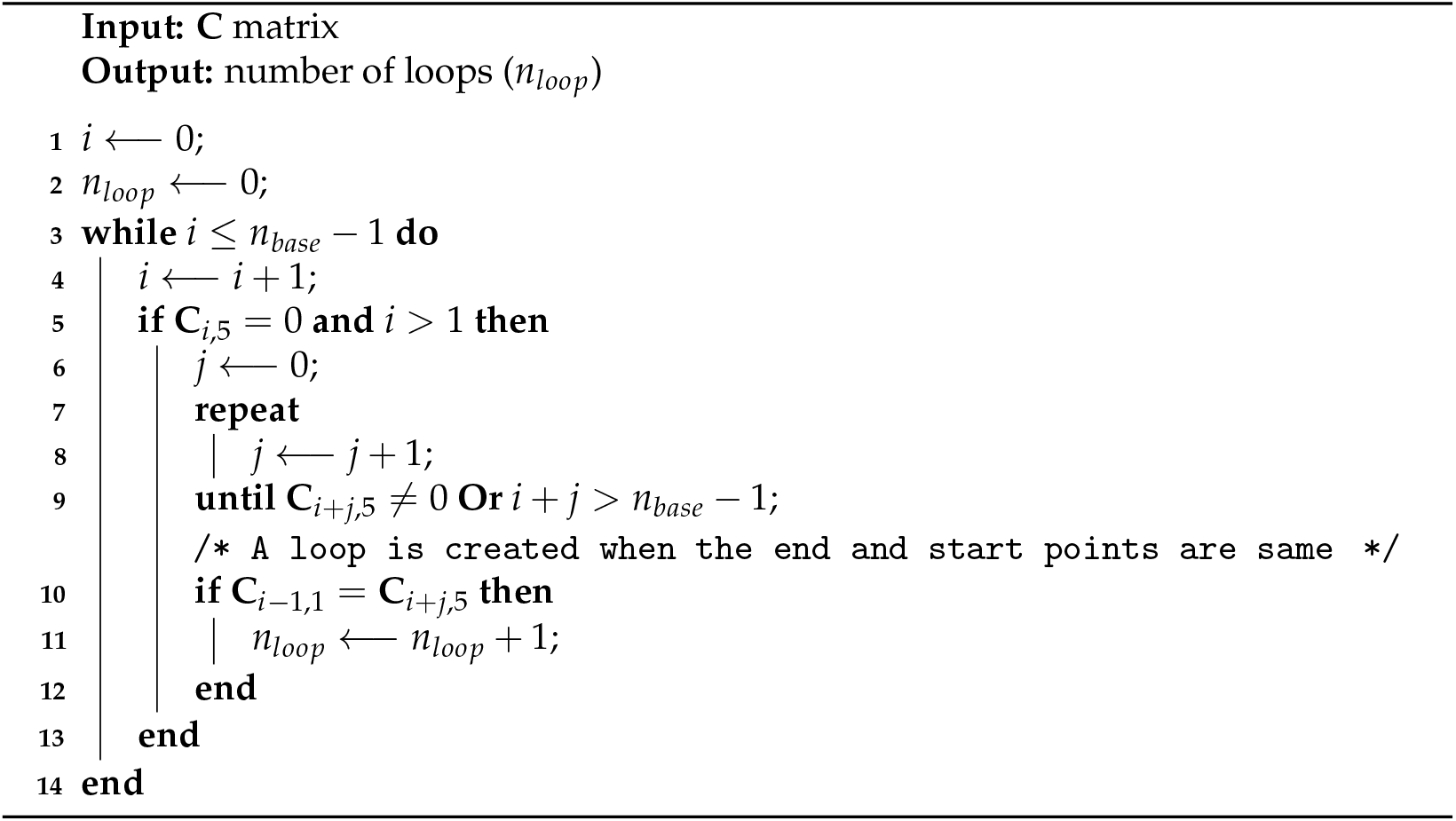

**Figure 2.**
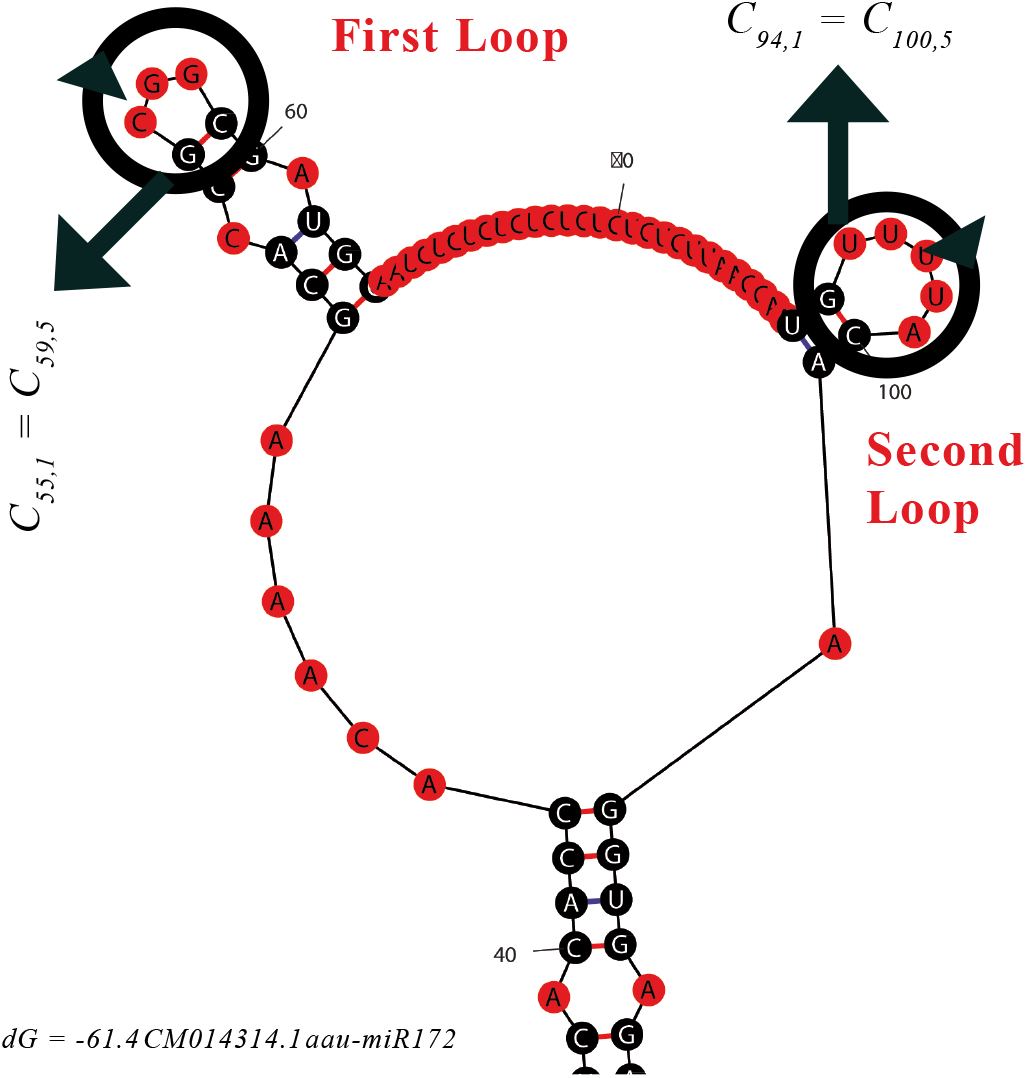
Demonstrating Algorithm 1 over *dG* = −61.4CM014314.1aau – miR172.

If the number of loops is more than one (i.e., multi-branched loop), *miRCurator* will mark the corresponding FASTA file with the red color for visual cues.

The second part of the *miRCurator* checks the gene for unacceptable structures. If the DICER cut point is located at exactly the mature strand start, it will be reported as unacceptable. Moreover, if the number of DICER cuts between the mature strand start and mature strand end is more than four then *miRCurator* will mark the corresponding gene as unacceptable. The reported genes in the second algorithm are painted in cyan color to provide visual cues for users who wants to double check the results. If a gene is filtered by both algorithms, it will be painted in magenta color (the combination of red and cyan). The time complexity of the second part is in order of 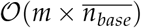. Figure 3 depicts the results of the second algorithm on a gene, in which the DICER cut is located at the mature strand start.

**Figure 3.**
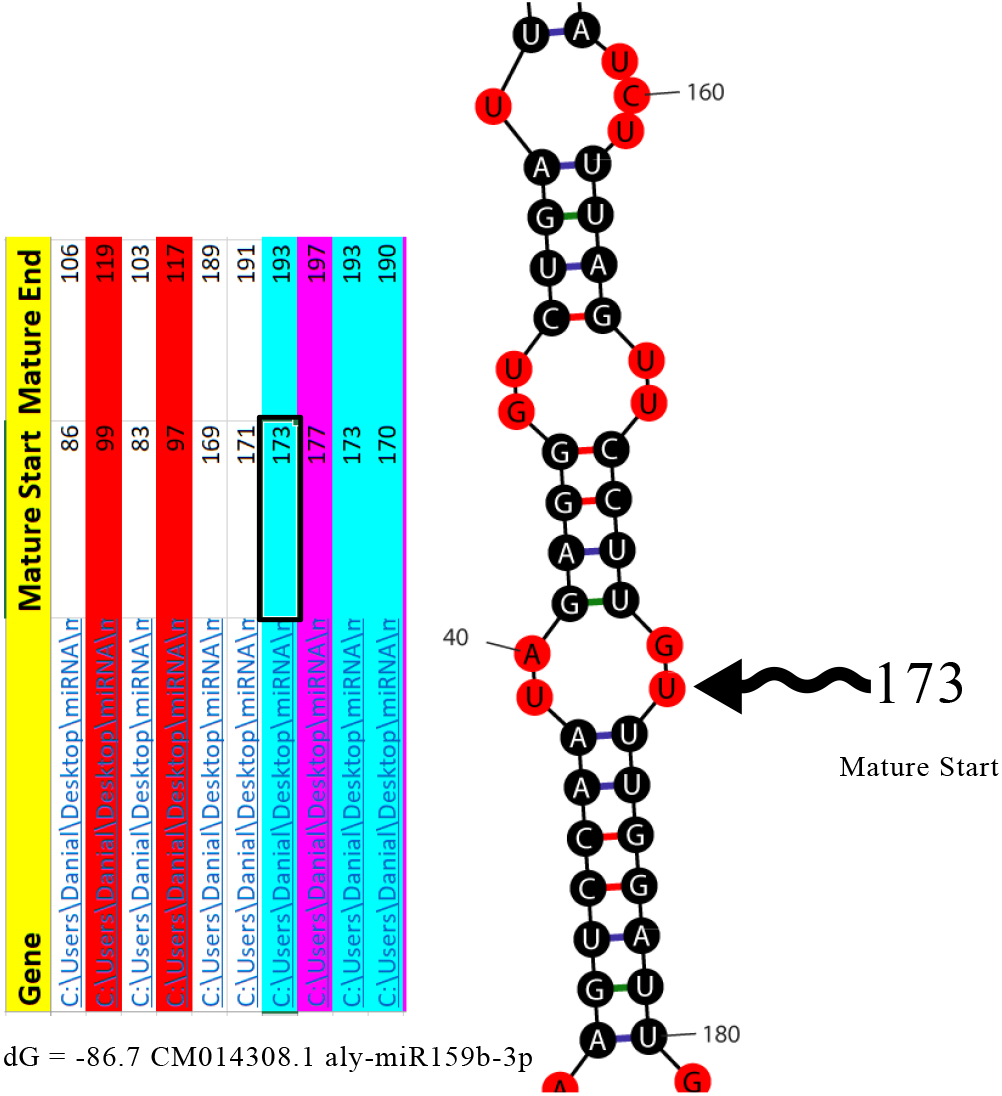
Illustrating a sample, in which the DICER cut coincides with the mature strand start.

Using a 64-bit computer that benefits from an Intel Core i5-4200U CPU and 4 GB memory, the developed script processed 4970 genes in 77.16 seconds. According to the MTM-1 method [18] and experts’ opinions, a manual review of such a dataset can take over a day.

After processing the whole dataset, *miRCurator* asks the user whether s/he is interested in a visual summary. The script generates a visual representation of results, which can be loaded with *Academic Presenter* [19] if the user approves. Due to its advantages and flexibility, *Academic Presenter* is being widely used for data visualization purposes [12,20]. As illustrated in Figure 4, the generated graph is a vector-based image that classifies the members into three categories: Group *A* represents those genes that are painted in cyan in the MS Excel file, Group *B* includes those that are eliminated because of multi-branched loops (marked with red color in the spreadsheet), and the magenta ones, at the center, are rejected by both algorithms (*A* ∩ *B*).

**Figure 4.**
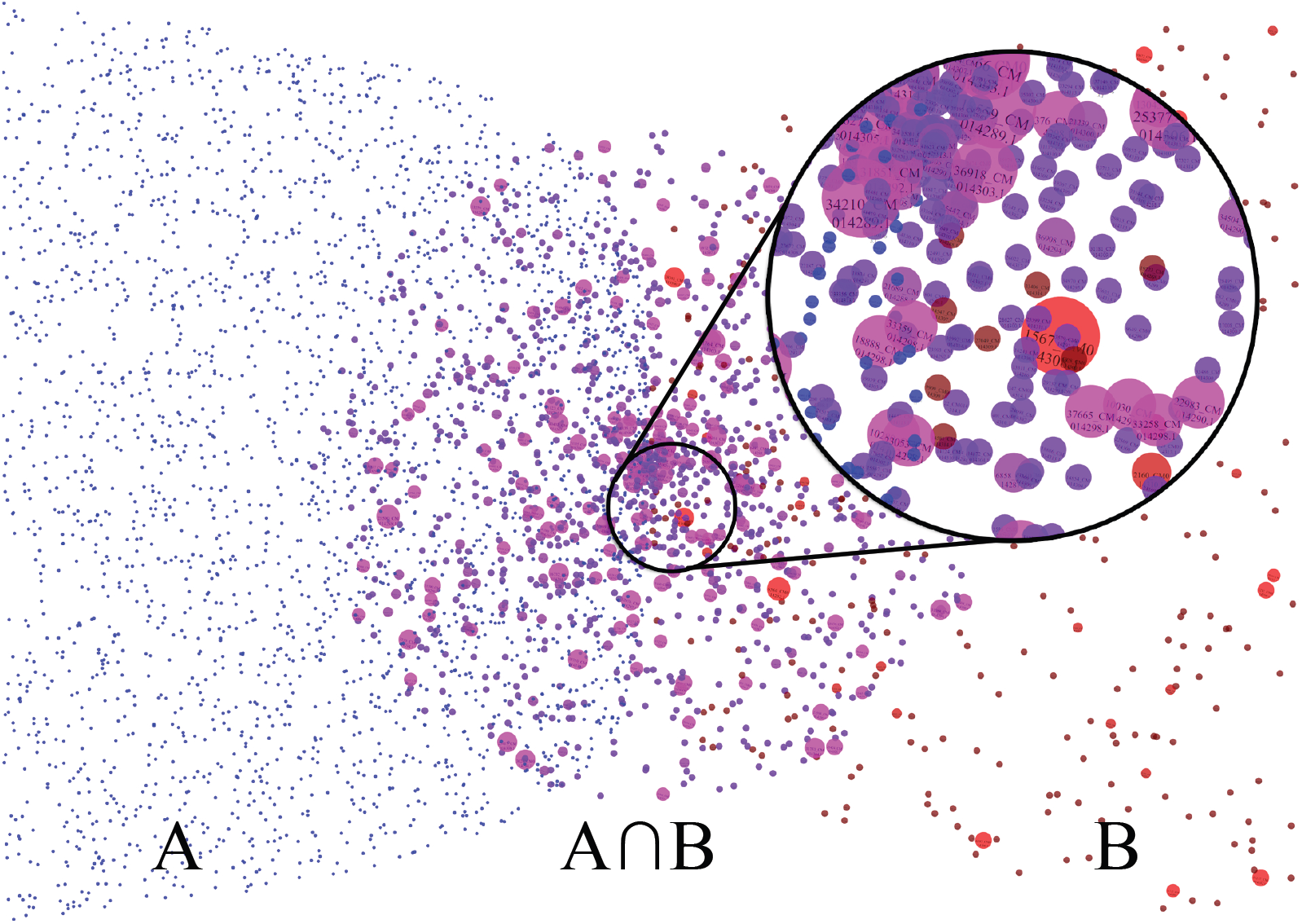
Generated visual for a sample dataset. As the number of branches increases in a gene, the radius of the corresponding blob grows and the color shifts toward red.

## 3. Results and Discussion

In all the studies mentioned in Section 2, the miRNA genes are identified by SUmiRFind and then, SUmiRFold script is used to decide the secondary structures by folding them. Using *miRCurator*, we successfully identified the pre-miRNAs, which have hairpins with multi-branched loops, inappropriate DICER cut sites, and more than four mismatches on their mature strand. Table 1 reports the number of pre-miRNAs, and the remaining pre-miRNAs after the filtration with *miRCurator* in detail for ten datasets. On average, *miRCurator* eliminates around 40.4% of ill-structured pre-miRNAs. To ameliorate the error in the calculations, the CPU time is computed as the average of three runs in each case study.

**Table 1:**
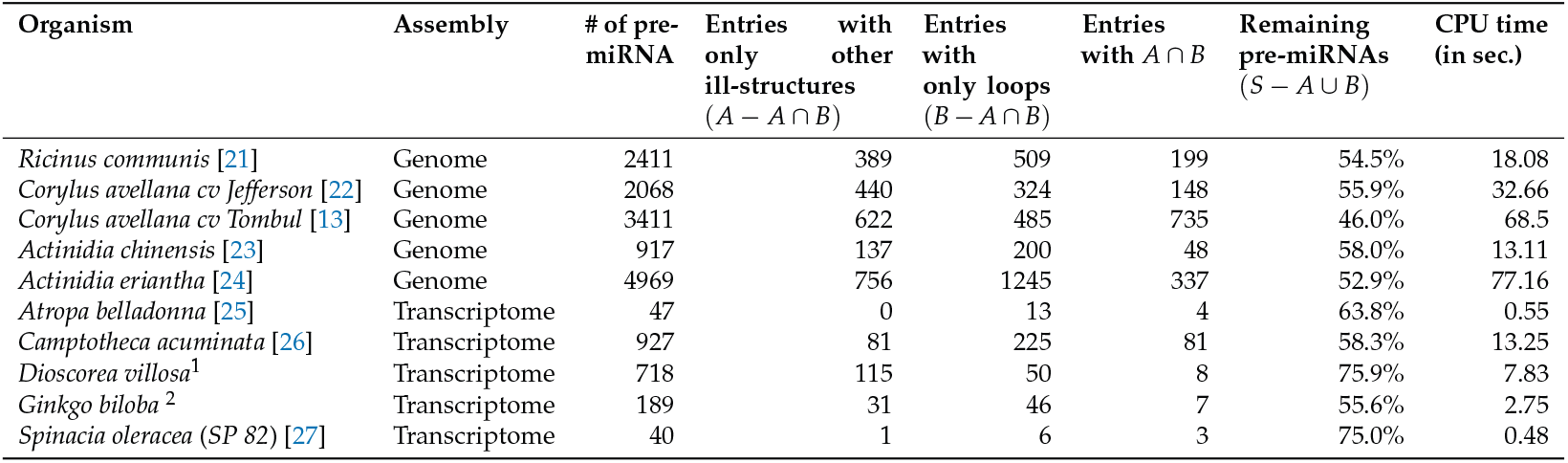
The summary of *miRCurator’s* output in various datasets from previously published studies.

We also conduct regression together with analysis of variance (ANOVA) on results in Table 1 to quantify the correlation between the CPU time (*y*) and regressors, which are the number of ill-structured entries (*x_A_* = |*A*|) and entries with multi-branched loops (*x_B_* = |*B*|). To prevent over-fitting, we set the constant to zero (*y* = *β*_1_*x_A_* + *β*_2_*x_B_*)^3^. The results show that there is a strong correlation between regressors and CPU time (*R*^2^ = 0.97). Additionally, the ANOVA test rejects H_0_-hypothesis, in which the coefficients of regressors are presumed zero (*β*_1_ = *β*_2_ = 0). Our analysis shows that the CPU time is more affected by *x_B_* (*β*_2_ = 0.028) than *x_A_* (*β*_1_ = 0.024). Figure 5 depicts the difference between the predicted and observed CPU time with respect to *x_A_* and *x_B_*.

**Figure 5.**
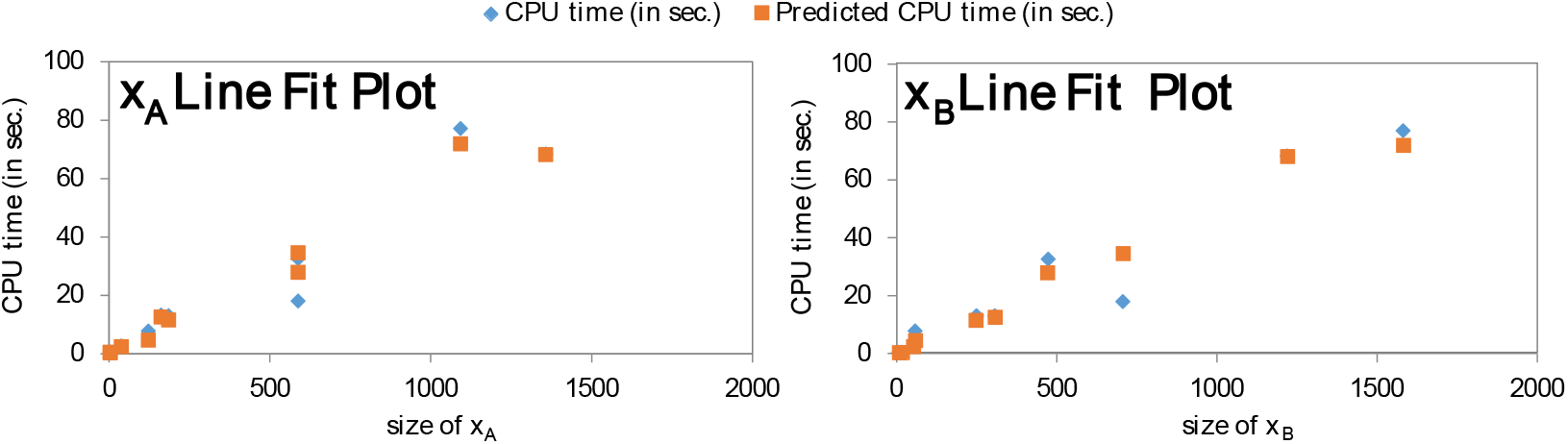
Predicted CPU time *ŷ* and observed CPU time *y* versus *x_A_* and *x_B_*.

Compared to the other Linux-based tools in the field [28], *miRCurator* is quick, accurate, and user-friendly since it provides outputs in MS Excel file format. In addition to these, our tool creates a detailed report at the end of the process that depicts all types of unwanted structures with their identification numbers. In the generated report, the users can perceive the information about the number of the multi-branched loops in each pre-miRNAs (the size of the red blobs), the number of high mismatch containing mature miRNAs and the number of the mature miRNAs having inappropriate DICER-cut points (blue-colored blobs). Although the generated MS Excel file by *miRCurator* with restricted color options provide visual cues, *Academic Presenter’s* image provide additional information using more color options, by which the users can identify the number of loops in pre-miRNAs. As a result of this, the users can conveniently track the various number of multi-branched formations.

The users may benefit from these outputs as the different number of multi-branched loops in a miRNA can be important to calculate the free energies of the secondary structures [29]. We also show the potential genes in the report, which are the combination of the multibranched loops and mismatches. By having all these outputs, the users can investigate the miRNA genes in detail for further studies.

## Author Contributions

Conceptualization, methodology, validation, data curation, formal analysis, B.A.; software, visualization, D.E.A.; writing—original draft preparation, B.A. and D.E.A.;

## Institutional Review Board Statement

Not applicable.

## Informed Consent Statement

Not applicable.

## Data Availability Statement

The interested readers can acquire the VBA script and a small case study from the supplementary material.

## Abbreviations

The following abbreviations are used in this manuscript:

miRNA: microRNA
dsRNA: double-stranded RNA
hpRNA: hairpin structured RNA
DCL: DICER-like
AGO: Argonaute
tasiRNA: trans-acting small interfering RNA
RDRs: RNA-dependent RNA polymerases
RISC: RNA-induced silencing complex
VBA: Visual Basic for Applications
ANOVA: analysis of variance

## Footnotes

1 Downloaded from Medicinal Plant Genomics Resource http://mpgr.uga.edu/index.shtml

2 Downloaded from Medicinal Plant Genomics Resource http://mpgr.uga.edu/index.shtml

3 With this regression surface, we assume that datasets with no multi-branched loop or unwanted structure are empty.

## References

1. Guo, Q.; Liu, Q.; A Smith, N.; Liang, G.; Wang, M.B. RNA silencing in plants: mechanisms, technologies and applications in horticultural crops. Current Genomics 2016, 17, 476–489.

2. Wang, J.; Mei, J.; Ren, G. Plant microRNAs: biogenesis, homeostasis, and degradation. Frontiers in plant science 2019, 10, 360.

3. Guigon, I.; Legrand, S.; Berthelot, J.F.; Bini, S.; Lanselle, D.; Benmounah, M.; Touzet, H. miRkwood: a tool for the reliable identification of microRNAs in plant genomes. BMC genomics 2019, 20, 1–9.

4. An, J.; Lai, J.; Sajjanhar, A.; Lehman, M.L.; Nelson, C.C. miRPlant: an integrated tool for identification of plant miRNA from RNA sequencing data. BMC bioinformatics 2014, 15, 1–4.

5. Numnark, S.; Mhuantong, W.; Ingsriswang, S.; Wichadakul, D. C-mii: a tool for plant miRNA and target identification. BMC genomics. BioMed Central, 2012, Vol. 13, pp. 1–10.

6. Lucas, S.J.; Budak, H. Sorting the wheat from the chaff: identifying miRNAs in genomic survey sequences of Triticum aestivum chromosome 1AL. PLoS One 2012, 7, e40859.

7. Kozomara, A.; Griffiths-Jones, S. miRBase: annotating high confidence microRNAs using deep sequencing data. Nucleic acids research 2014, 42, D68–D73.

8. Camacho, C.; Coulouris, G.; Avagyan, V.; Ma, N.; Papadopoulos, J.; Bealer, K.; Madden, T.L. BLAST+: architecture and applications. BMC bioinformatics 2009, 10, 1–9.

9. Markham, N.R.; Zuker, M. UNAFold. In Bioinformatics; Springer, 2008; pp. 3–31.

10. Zhang, B.; Pan, X.; Wang, Q.; Cobb, G.P.; Anderson, T.A. Computational identification of microRNAs and their targets. Computational biology and chemistry 2006, 30, 395–407.

11. Avşar, B.; Esmaeili Aliabadi, D. Putative microRNA analysis of the kiwifruit Actinidia chinensis through genomic data. International Journal of Life Sciences Biotechnology and Pharma Research 2015, 4, 96–99.

12. Avşar, B.; Esmaeilialiabadi, D. Identification of microRNA elements from genomic data of European hazelnut (Corylus avellana L.) and its close relatives. Plant Omics 2017, 10, 190–196.

13. Lucas, S.J.; Kahraman, K.; Avşar, B.; Buggs, R.J.; Bilge, I. A chromosome-scale genome assembly of European hazel (Corylus avellana L.) reveals targets for crop improvement. The Plant Journal 2021, 105, 1413–1430.

14. Avşar, B.; Esmaeili Aliabadi, D. In silico analysis of microRNAs in Spinacia oleracea genome and transcriptome. International Journal of Bioscience, Biochemistry and Bioinformatics 2017, 7, 84–92.

15. Avsar, B.; Esmaeili Aliabadi, D. Identification of in silico mirnas in four plant species from fabaceae family. Agrofor 2018, 3.

16. Avsar, B.; Zhao, Y.; Li, W.; Lukiw, W.J. Atropa belladonna expresses a microRNA (aba-miRNA-9497) highly homologous to Homo sapiens miRNA-378 (hsa-miRNA-378); both miRNAs target the 3’-Untranslated region (3’-UTR) of the mRNA encoding the neurologically relevant, zinc-finger transcription factor ZNF-691. Cellular and molecular neurobiology 2020, 40, 179–188.

17. Avşar, B.; Aliabadi, D. In silico identification of microRNAs in 13 medicinal plants. Turkish Journal of Biochemistry 2018, 42, 57.

18. Karger, D.W.; Hancock, W.M. Advanced work measurement; Industrial Press, 1982.

19. Avşar, B.; Aliabadi, D.E.; Aliabadi, E.E.; Yousefnezhad, R. Academic Presenter: A new storytelling presentation software for academic purposes. arXiv preprint arXiv:1607.06979 2016.

20. Aliabadi, D.E.; Avşar, B.; Yousefnezhad, R.; Aliabadi, E.E. Investigating global language networks using Google search queries. Expert Systems with Applications 2019, 121, 66–77.

21. Chan, A.P.; Crabtree, J.; Zhao, Q.; Lorenzi, H.; Orvis, J.; Puiu, D.; Melake-Berhan, A.; Jones, K.M.; Redman, J.; Chen, G.; others. Draft genome sequence of the oilseed species Ricinus communis. Nature biotechnology 2010, 28, 951–956.

22. Rowley, E.R.; VanBuren, R.; Bryant, D.W.; Priest, H.D.; Mehlenbacher, S.A.; Mockler, T.C. A draft genome and high-density genetic map of European hazelnut (Corylus avellana L.). BioRXiv 2018, p. 469015.

23. Huang, S.; Ding, J.; Deng, D.; Tang, W.; Sun, H.; Liu, D.; Zhang, L.; Niu, X.; Zhang, X.; Meng, M.; others. Draft genome of the kiwifruit Actinidia chinensis. Nature communications 2013, 4, 1–9.

24. Tang, W.; Sun, X.; Yue, J.; Tang, X.; Jiao, C.; Yang, Y.; Niu, X.; Miao, M.; Zhang, D.; Huang, S.; others. Chromosome-scale genome assembly of kiwifruit Actinidia eriantha with single-molecule sequencing and chromatin interaction mapping. GigaScience 2019, 8, giz027.

25. Bedewitz, M.A.; Góngora-Castillo, E.; Uebler, J.B.; Gonzales-Vigil, E.; Wiegert-Rininger, K.E.; Childs, K.L.; Hamilton, J.P.; Vaillancourt, B.; Yeo, Y.S.; Chappell, J.; others. A root-expressed L-phenylalanine: 4-hydroxyphenylpyruvate aminotransferase is required for tropane alkaloid biosynthesis in Atropa belladonna. The Plant Cell 2014, 26, 3745–3762.

26. Zhao, D.; Hamilton, J.P.; Pham, G.M.; Crisovan, E.; Wiegert-Rininger, K.; Vaillancourt, B.; DellaPenna, D.; Buell, C.R. De novo genome assembly of Camptotheca acuminata, a natural source of the anti-cancer compound camptothecin. GigaScience 2017, 6, gix065.

27. Xu, C.; Jiao, C.; Zheng, Y.; Sun, H.; Liu, W.; Cai, X.; Wang, X.; Liu, S.; Xu, Y.; Mou, B.; others. De novo and comparative transcriptome analysis of cultivated and wild spinach. Scientific reports 2015, 5, 1–9.

28. Alptekin, B.; Akpinar, B.A.; Budak, H. A comprehensive prescription for plant miRNA identification. Frontiers in plant science 2017, 7, 2058.

29. Diamond, J.M.; Turner, D.H.; Mathews, D.H. Thermodynamics of three-way multibranch loops in RNA. Biochemistry 2001, 40, 6971–6981.

